# Sensitization of Chrysosplenetin against Artemisinin Resistant *Plasmodium berghei* K173 via Blocking Host ABC Transporters Potentially Mediated by NF-κB p52 or PXR/CAR Signaling Pathway

**DOI:** 10.1101/2022.01.10.475770

**Authors:** Xuesong Zhao, Shanhong Ni, Yuanyuan Zhang, Xiuli Wu, Ying Yao, Jing Chen

## Abstract

This study investigated if artemisinin-chrysosplenetin combination (ART-CHR) improved ART antimalarial efficacy against resistant *Plasmodium berghei* K173 *via* depressing host ABC transporter and potential molecular mechanism. Parasitaemia% and inhibition% were calculated and gene/protein expressions of ABC transporters or PXR/CAR/NF-κB p52 were detected by Western-blot and RT-qPCR. *In vitro* transcription of PXR/CAR was studied by dual-luciferase reporter assay. Our data indicated that ART-CHR improved ART efficacy against resistant parasites. P-gp inhibitor verapamil and CHR showed a stronger effect in killing resistant parasites while vehicle and Bcrp inhibitor novobiocin did not. ART activated intestinal ABCB1/ABCG2 and CHR inhibited them. ART decreased Bcrp protein whereas CHR increased it. ART ascended ABCC1/ABCC4/ABCC5 mRNA but ART-CHR descended them. CHR as well as rifampin (RIF) or 5-fluorouracil (5-FU) increased transcription levels of PXR/CAR while showed a versatile regulation on *in vivo* hepatic and enternal PXR/CAR in Mdr1a+/+ (WT) or Mdr1a-/- (KO) mice infected with sensitive or resistant parasites. Oppositely, hepatic and enteric N-κB p52 mRNA was conformably decreased in WT but increased in KO-resistant mice. NF-κB pathway should potentially involved in the mechanism of CHR on inhibiting ABC transporters and ART resistance while PXR/CAR play a more complicated role in this mechanism.

## INTRODUCTION

Artemisinin-Based Combination Treatments (ACTs) are the utmost important worldwide combination therapy of *Plasmodium falciparum* malaria (1-2). Unfortunately, ARTs resistance, defined as a delayed clearance of parasites, has clinically occurred (3) possessing undefined mechanisms where ABC transporter super-family might involve (4). P-glycoprotein (P-gp, gene symbol ABCB1 or MDR1), multi-drug resistance protein 1-6 (MRP1-6, gene symbol ABCC1-6), and breast cancer resistance protein (BCRP, gene symbol ABCG2) are therein the best distinguished transporters directly leading to MDR (5-7). ARTs induced CYP2B6, 3A4, and MDR1 expressions *via* activating transcription levels of human pregnane X receptor (hPXR) and human/mouse constitutive androstane receptor (hCAR or mCAR) (5). Meanwhile, activated nuclear factor kappa-B (NF-κB) promotes over 150 target transcripts signifying the potential of NF-κB signaling pathway on ABC transporters (8-9).

Polymethoxylated flavonoids are discovered to modulate phase I CYP450s or phase II UGTs and ABC transporters, which potentially alter pharmaceutics of substrate drugs (10-12). Chrysosplenetin (CHR) is one of known polymethoxylated flavonoids in leaf of *Artemisia annua* L. (13). Previously, CHR purified from ART industrial waste improved the bioavailability and anti-malarial efficiency of ART in combination ratio of 1:2 (optimized from combination proportions of 1:0.5, 1:1, 1:2, and 1:4 between ART and CHR) through strongly inhibiting rat CYP3a activity in an un-competitive manner (14) and *in vivo* or *in vitro* P-gp efflux (15-16) *via* reversing the up-regulated P-gp/ABCB1 or ABCG2 mRNA levels by ART in healthy mice (16-17). Here, we further design this experiment to explicit the sensitization of CHR against ART resistant *Plasmodium berghei* K173 by blocking host ABC transporters and upstream transcript regulations by PXR/CAR and NF-κB k52.

## RESULTS

### Erythrocyte entry by sensitive and resistant malarial parasites

Parasitaemia% at different experimental time periods (0-180 h) indicated different infection curves in erythrocytic stage between sensitive and resistant parasites (Figure 1). Within 100 h, the two types of parasites remained similarly slow endoerythrocytic infection rate but parasitaemia% of the sensitives increased steeply after that point of time. Resistant parasites, however, kept lower parasitaemia%, requiring longer infection time than the sensitives.

**Fig. 1.**
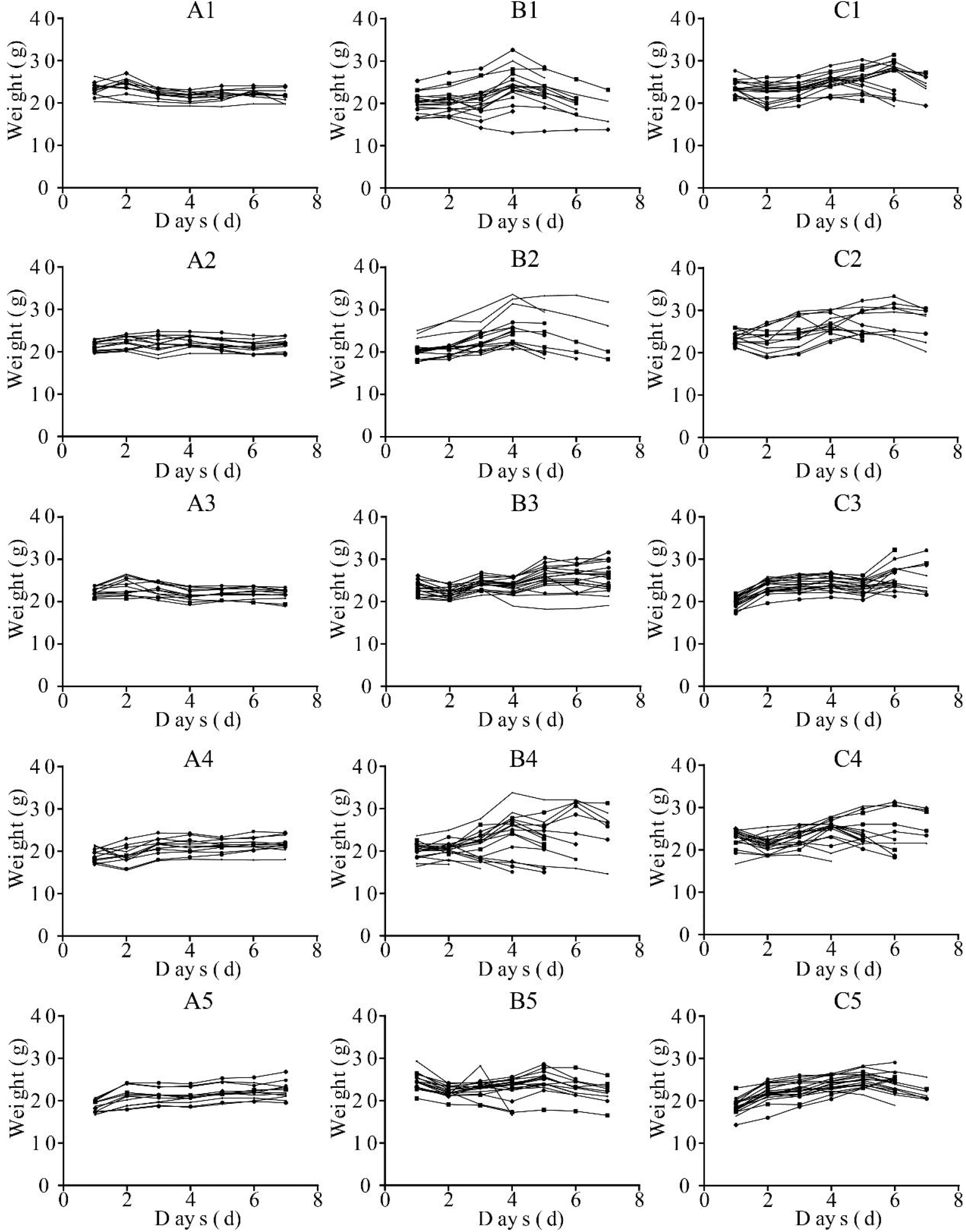
Body weights for healthy (A1-A5) and ART-sensitive (B1-B5) or -resistant (C1-C5) malaria parasites infected mice. Mice in the A1, B1, and C1 groups were treated by CMC-Na. A2, B2, and C2 groups were treated by VER alone. A3, B3, and C3 groups were treated by ART alone. A4, B4, and C4 groups were treated by CHR alone. A5, B5, and C5 groups were treated by ART-CHR combination.

The morphology of infected RBC was described in Figure 2A1-A5, B1-B5, and C1-C5. It is observable, pre- and post-medication, that ART-resistant *P. berghei* in erythrocytic stage experienced a developmental retardation versus the sensitives. RBC infected with sensitive parasites became deformed severely while those with resistant parasites remains normal shape.

**Fig. 2.**
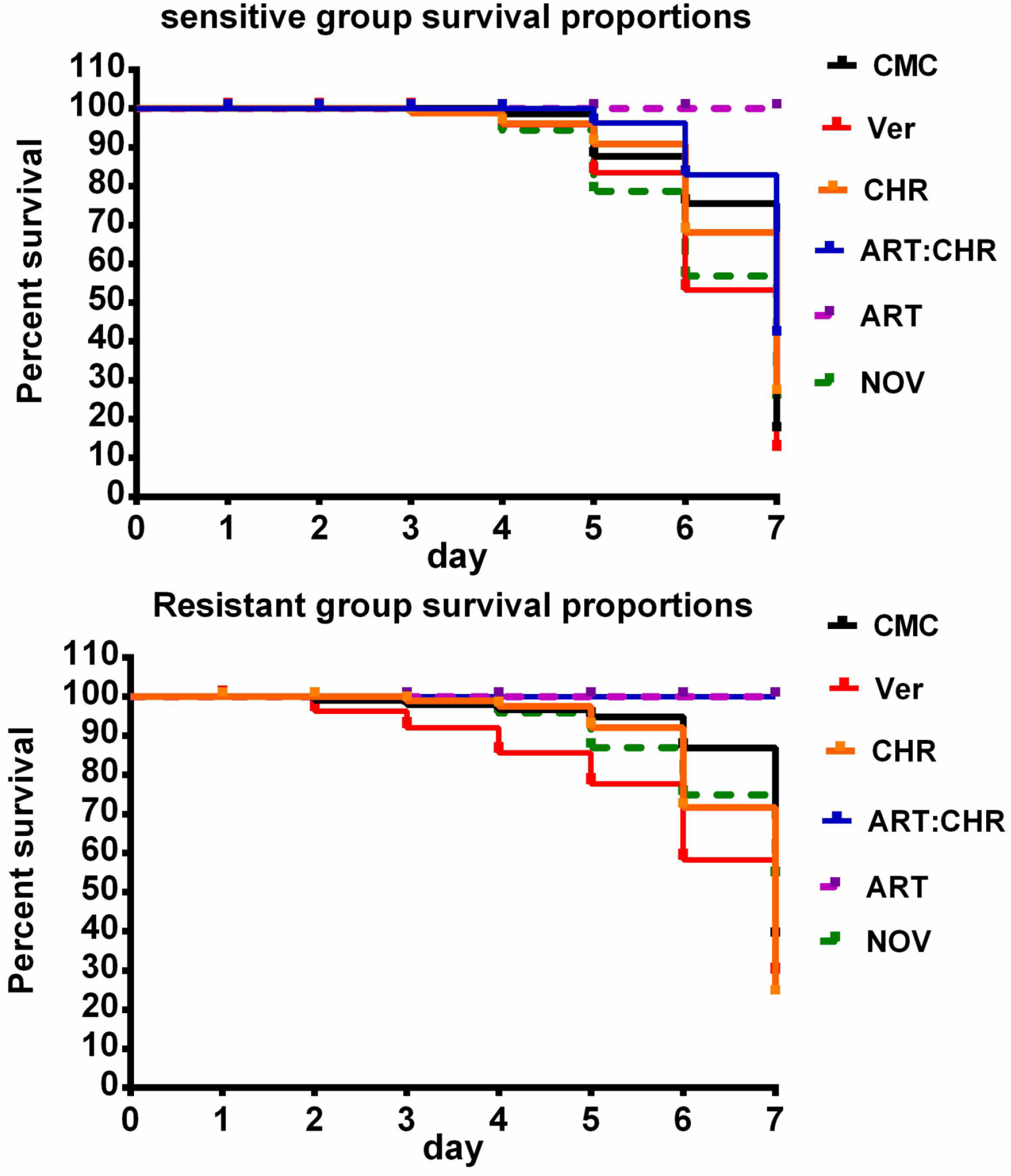
Kaplan-Meier mice survival curves statistically using Log-rank (Mantel-Cox) test (Graphpad 6.0, USA) after being infected with ART-sensitive (A) or -resistant malaria (B) parasites under different drug treatments.

### Body weights and survivals

Body weight curves were displayed in Figure 3A1-A5, B1-B5, and C1-C5. Unlike the stabilized body weights of healthy mice, the curves for the mice infected by ART-sensitive or resistant parasites fluctuated while ART-sensitive *P. berghei* infection led to more turbulent weight gain of mice. After treated by ART or ART-CHR combination, it became normalized. Both VER and CHR stabilized body weights in resistant groups despite no changes occurred in sensitives.

**Fig. 3.**
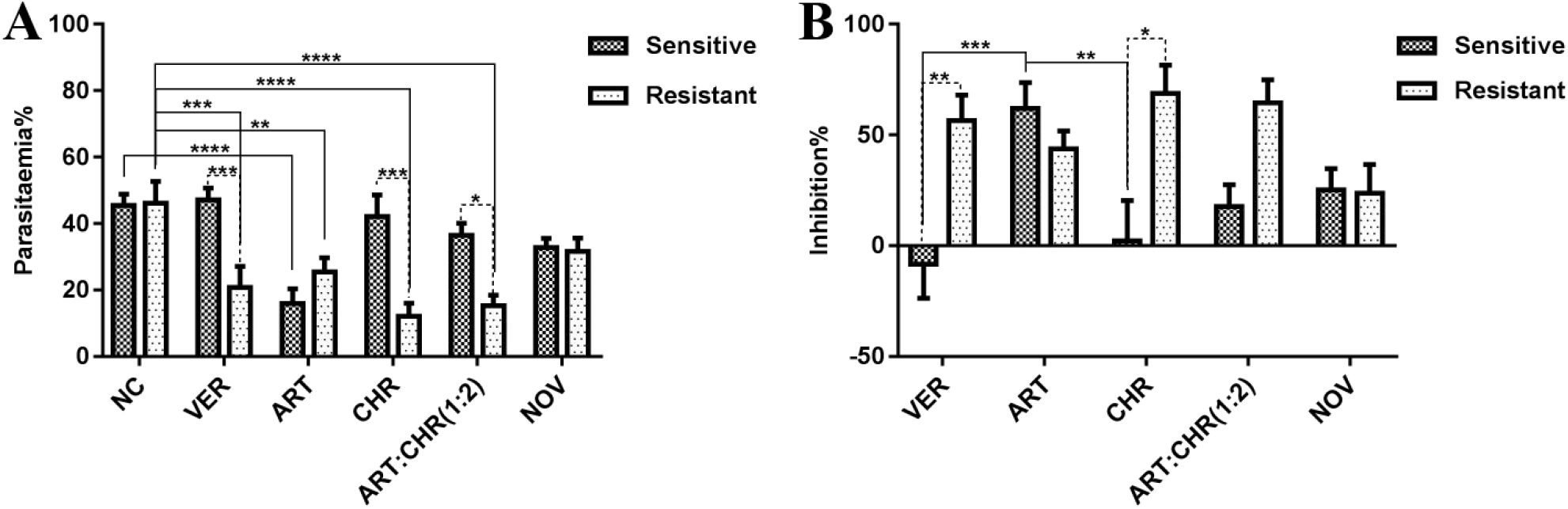
Parasitaemia% (A) and inhibition% (B) under different drug treatments.

Mice survivals were displayed in Figure 4A and 4B or Table 1. Slightly decreased percent survival from 100% to 94.44% was observed after 7 days of ART treatment against resistant *P. berghei* infected mice versus sensitive groups, indicating a potential risk of its resistance. When treated by ART-CHR combination, mice in the resistant group all survived while survival of sensitive group was only 50.67%.

**TABLE 1.**
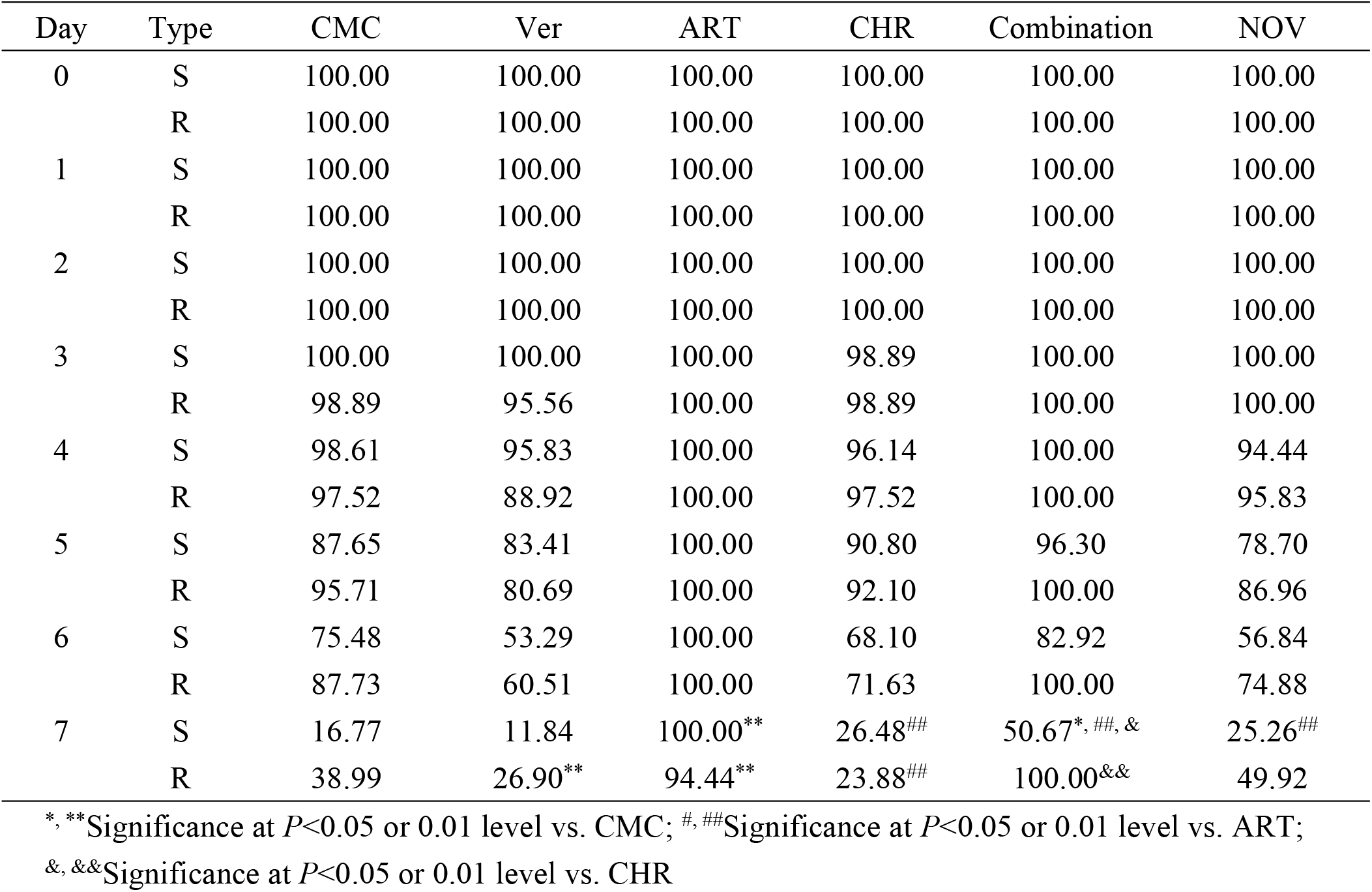
The Kaplan-Meier analysis in ART-sensitive (S) or -resistant groups (R) under different drug treatments.

**Fig. 4.**
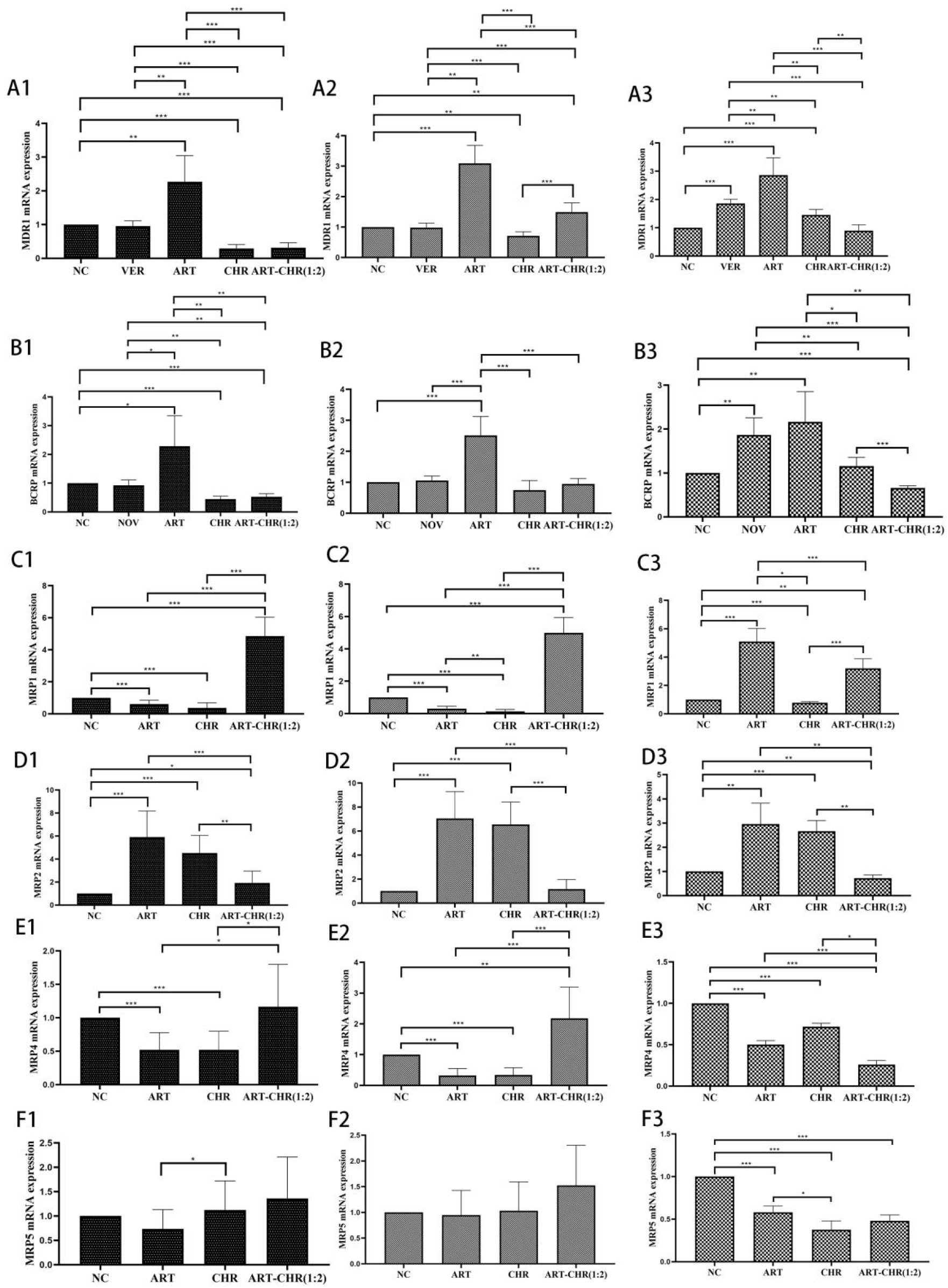
ABC transporter mRNA levels were detected by real-time qPCR after 7 days of drug treatments in host small intestines from healthy (A1-F1), ART-sensitive (A2-F2), and -resistant malaria parasites infected ICR mice (A3-F3). CMC-Na was used as negative control and VER or NOV as P-gp or Bcrp positive inhibitors, respectively. All values were expressed as the mean ± S.D. (*n*=6).

### Anti-malarial activity of ART with or without CHR against ART-sensitive and - resistant parasites

In Figure 5A and 5B, NOV had no anti-malarial efficacy and did not lead to a different parasitaemia% or inhibition% between two groups, similarly to NC. ART decreased parasitaemia% and showed a strong killing effect against sensitive parasites versus NC but its efficacy against resistant *P. berghei* slightly declined. Intriguingly, VER, CHR, and ART-CHR combination decreased parasitaemia% and increased inhibition% at least 2 folds against ART-resistant *P. berghei* than NC or the sensitives. They killed resistant parasites but less so in sensitive groups.

**Fig. 5.**
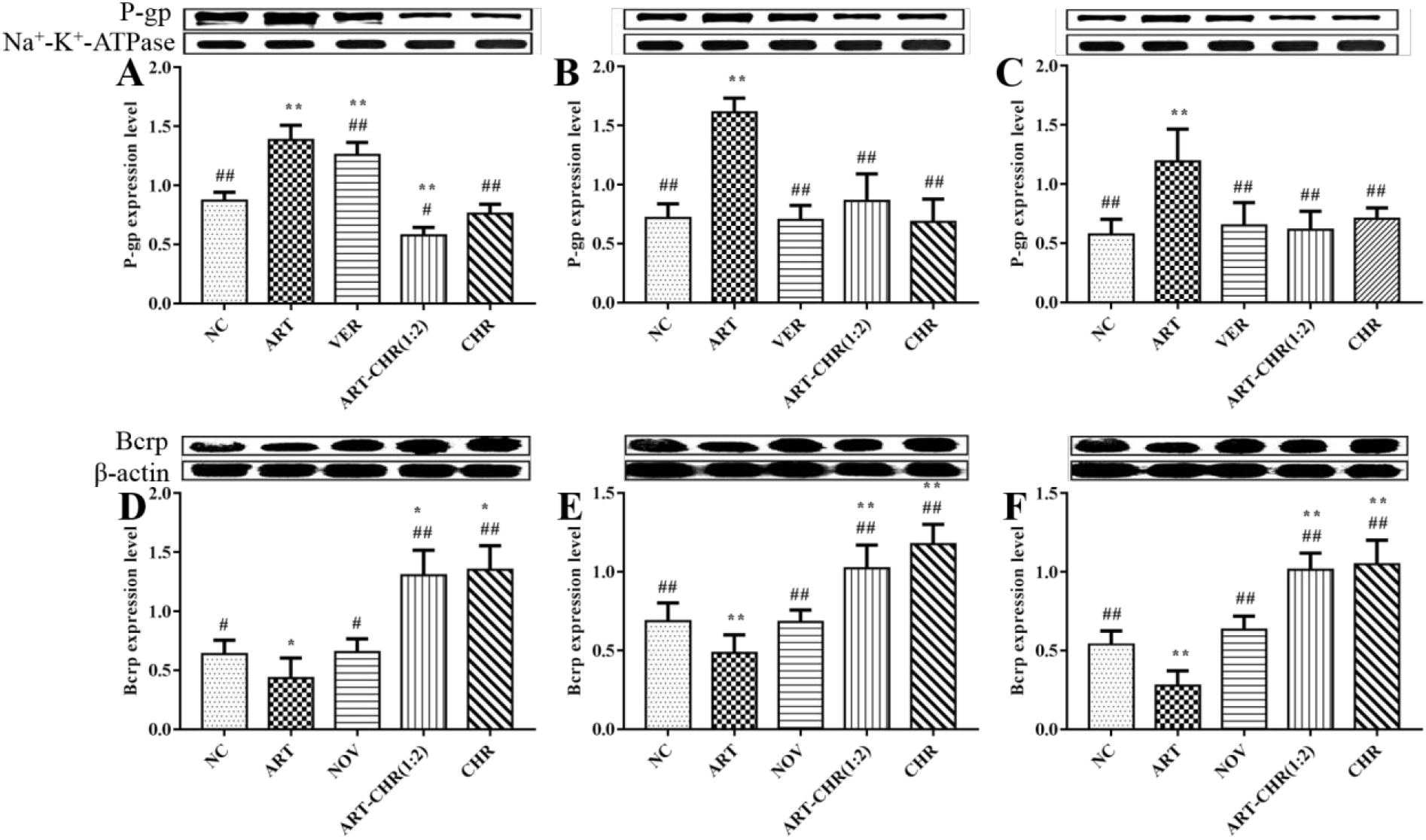
P-gp/Bcrp expression levels determined by western-blot analysis for healthy (A and D), ART-sensitive (B and E), and -resistant (C and F) malaria parasites infected mice. Quantification of P-gp/Bcrp assessed by western blotting analysis was normalized to the expression level of Na^+^-K^+^-ATPase/β-actin antibody. All values were expressed as the mean ± S.D. (*n*=6). \**P*<0.05 and \*\**P*<0.01 vs negative control. ^#^*P*<0.05 and ^##^ *P*<0.01 vs ART alone.

### Gene expressions of ABC transporters in small intestines

Data indicated that ART up-regulated ABCB1/ABCG2 mRNA expressions in small intestines from both healthy and parasites infected mice versus NC. CHR and ART-CHR combination down-regulated them before and after wild or mutant parasite infection (Figure 6A1-A3 and B1-B3). VER and NOV oppositely increased ABCB1/ABCG2 mRNA expressions in resistant groups despite no changes in healthy or sensitive groups.

**Fig. 6.**
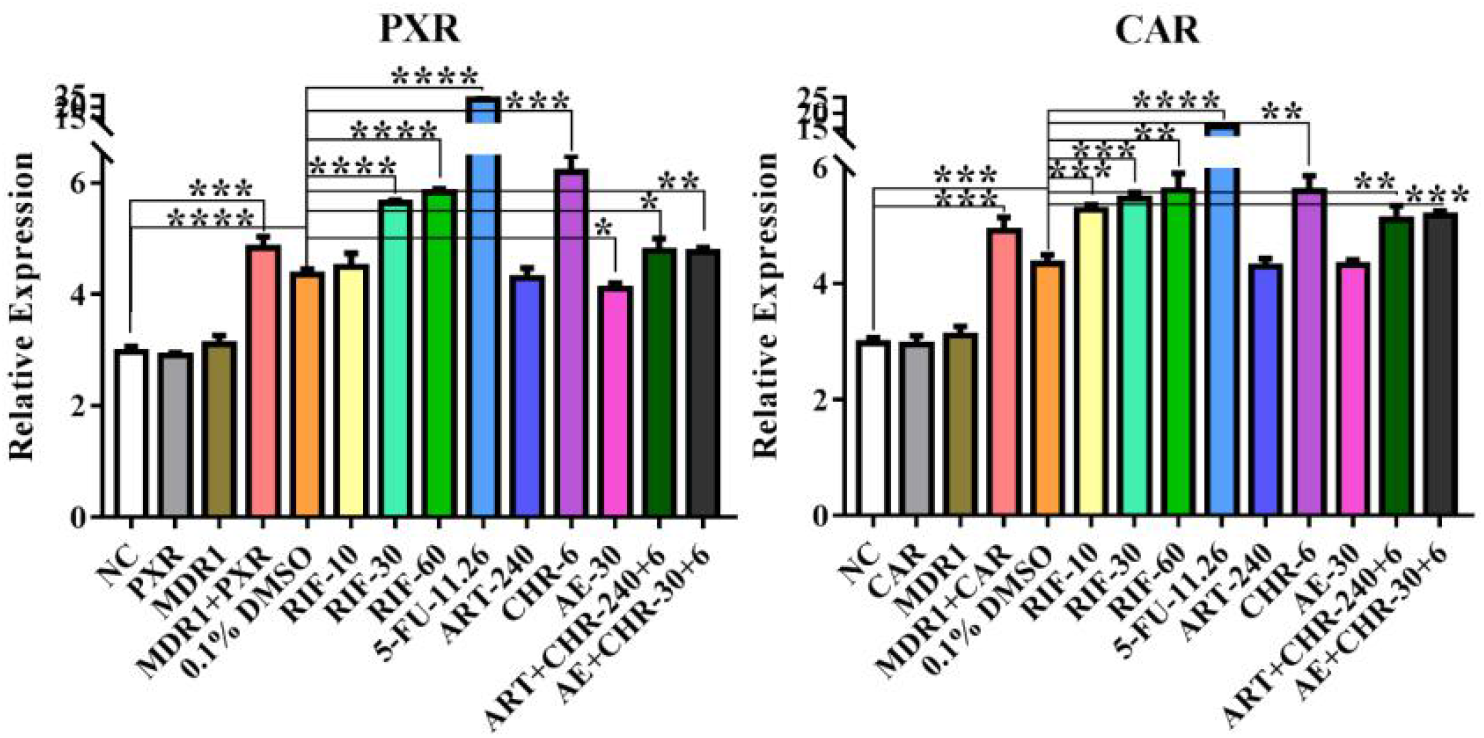
Impact of different drugs on the transcriptional expression of hPXR/hCAR detected by a dual-luciferase reporter assay in the human intestinal cell lines LS174T. All values were expressed as the mean ± S.D. (*n*=6). \**P*<0.05 and \*\**P*<0.01 vs negative control.

ART decreased MRP1 mRNA in healthy or sensitive mice whereas increased it in resistant groups versus NC. CHR descended it in mice pre- and post-infection with wild and mutant parasites. ART-CHR ascended MRP1 mRNA in healthy and sensitive groups but decreased it in resistant groups (Figure 6C1-C3). ABCC2 mRNA was up-regulated by ART or CHR while down-regulated by ART-CHR combination (Figure 6D1-D3). ART and CHR decreased ABCC4 mRNA which was enhanced by combination in healthy and sensitive groups but declined in resistant groups (Figure 6E1-E3). Similarly, combination heightened ABCC5 mRNA level in healthy and sensitive groups whereas descended it in resistant groups (Figure 6F1-F3).

### Protein expression of P-gp and Bcrp in small intestines infected with ART-sensitive and -resistant *P. berghei*

As shown in Figure 7A-F, completely opposite results were discovered. ART facilitated P-gp expression while suppressed Bcrp protein levels, whether the mice were infected by parasites or not. VER led to an increased P-gp expression in healthy mice when compared with NC but a lower level in sensitive or resistant *P. berghei* infected mice versus ART. CHR and ART-CHR combination consistently down-regulated P-gp but activated Bcrp expression versus NC and ART in healthy and parasite-infected mice.

**Fig. 7.**
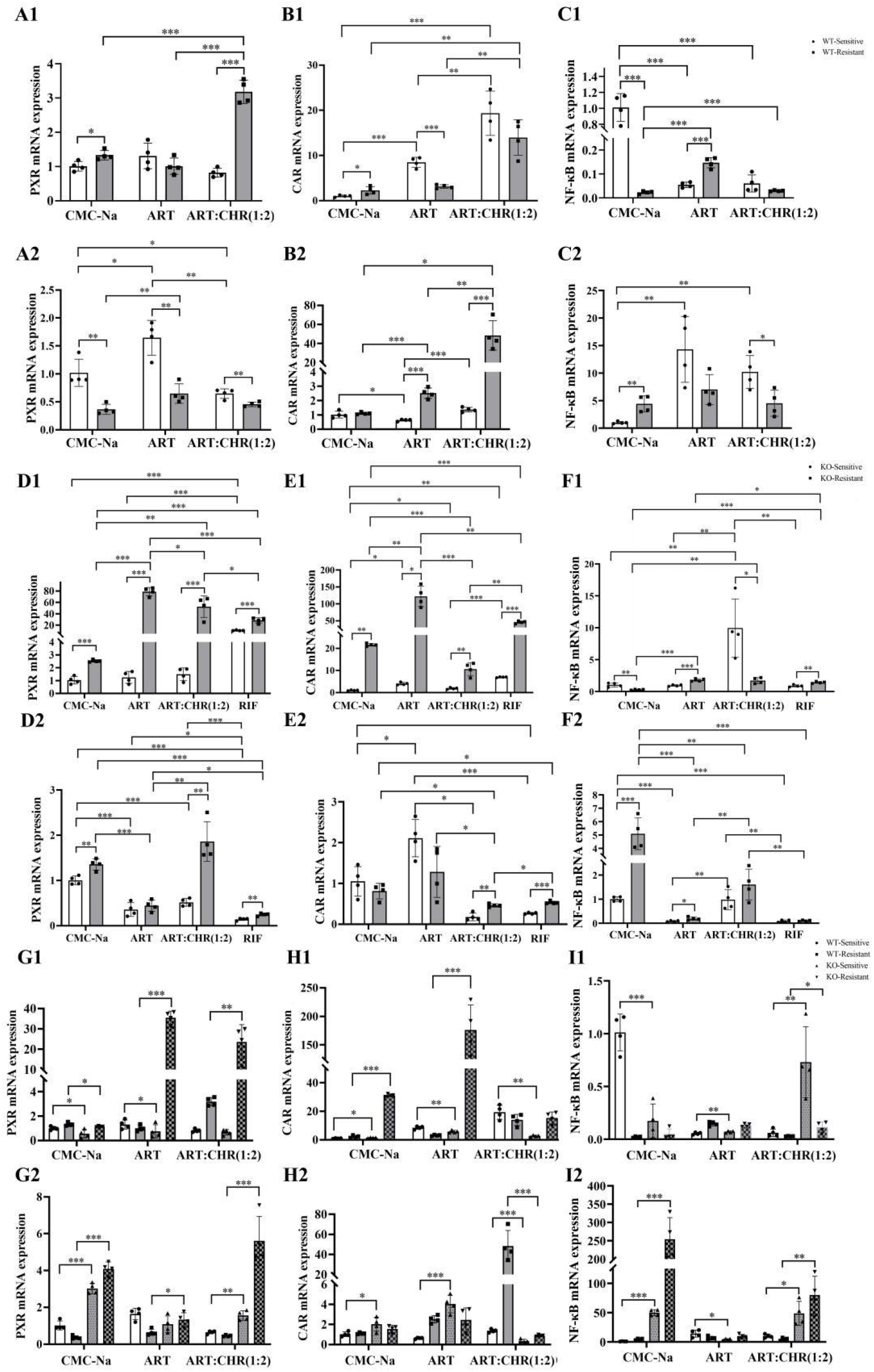
The PXR/CAR and NF-κB mRNA levels detected by real-time qPCR (group A, B, and C: WT-sensitive or resistant; A1-C1: livers; A2-C2: small intestines; group D, E, and F: KO-sensitive or resistant; D1-F1: livers; D2-F2: small intestines; group G, H, and I: WT-sensitive or resistant and KO-sensitive or resistant; G1-I1: livers; G2-I2: small intestines). WT and KO respectively represented wide type and gene deficiency of Mdr1a, encoding gene of P-gp. CMC-Na was used as negative control and rifampicin (RIF) as a P-gp inducer, positive control. All values were expressed as the mean ± S.D. (*n*=4).

### Transcriptional expressions of PXR and CAR in LS174T cell line impacted by ART with or without CHR

Firstly, IC_50_ values of 5-FU, CHR, and ART in LS174T were measured to be 9.30 ± 0.73, 4.17 ± 0.74, and 153.73 ± 17.81 μM, providing the appropriate concentrations for administration. Results indicated that the relative expression level of MDR1 or PXR has no difference when they were separately transfected to the plasmid versus NC, while it was increased by 1.6 (in DMEM only) and 1.5 folds (in DMEM containing 0.1%DMSO) after a transient co-transfection of MDR1 and PXR, indicating that PXR, as the upstream gene regulator, activated MDR1 expression (Figure 8). The similar result was observed in dual-luciferase reporter system for CAR. Therefore, dual-luciferase reporter system for PXR/CAR was successfully constructed.

RIF, a positive drug, at dosages of 10, 30, and 60 μM significantly increased PXR and CAR relative levels. 5-FU also remarkably activated PXR and CAR transcriptional expressions. CHR and ART-CHR combination slightly increased PXR and CAR levels while ART did not lead to a difference compared with NC which is not in accordance with the literature (5). AE slightly down-regulated PXR but in combination with CHR both PXR and CAR were increased.

### *In vivo* PXR/CAR and NF-κB p52 levels in livers and small intestines of Mdr1a WT or KO mice after being infected with sensitive and resistant parasites under ART treatment with or without CHR

Firstly, in WT-sensitive or resistant groups (Figure 9A1-C1 and A2-C2), NC led to enhanced PXR/CAR mRNA and lessened NF-κB p52 mRNA in the livers but caused an decreased PXR along with increased NF-κB in small intestines from resistant groups than sensitives. In like manner, ART reduced hepatic CAR and enteral PXR or NF-κB while fortified intestinal CAR or hepatic NF-κB. Combination elevated hepatic PXR and intestinal CAR but lowered intestinal PXR or hepatic CAR as well as descended NF-κB in both livers and small intestines. In contrast with NC or ART, ART-CHR combination led to a higher hepatic PXR as well as a superior CAR in both livers and small intestines of resistant groups but lower NF-κB p52 mRNA (Figure 9A1, B1, B2, C1, and C2).

Secondly, in KO-sensitive or resistant groups (Figure 9D1-F1 and D2-F2), NC augmented both hepatic and enteral PXR mRNA as well as elevated hepatic CAR and enteric NF-κB whereas degraded hepatic NF-κB from resistant groups versus sensitives. Accordingly, ART increased hepatic PXR/CAR or NF-κB in both two issues; meanwhile, combination up-regulated liver and gut PXR/CAR mRNA while declined hepatic NF-κB accompanied with slightly increased intestinal NF-κB. RIF increased almost all the parameters except enteric NF-κB in resistant groups. When compared with NC, ART, ART-CHR, and RIF got a higher hepatic PXR mRNA in KO-resistant. ART contributed to a higher level of hepatic PXR in resistant groups than NC, ART-CHR, or RIF. ART-CHR also indicated a stronger induction of PXR than NC and RIF (Figure 9D1). ART and RIF increased hepatic CAR mRNA from KO-resistant while combination decreased it (Figure 9E1). ART-CHR activated NF-κB gene expression in livers from KO-sensitive and ART, ART-CHR, and RIF slightly induced hepatic NF-κB in KO-resistant versus NC (Figure 9F1). ART-CHR significantly enhanced enteral PXR mRNA in KO-resistant by comparison with NC, ART, and RIF (Figure 9D2). Both ART-CHR and RIF decreased enteric CAR in contrast with NC or ART (Figure 9E2). Intestinal NF-κB declined in both KO-sensitive and -resistant under ART and RIF treatments. ART-CHR led to a lower NF-κB in KO-resistant than NC and a higher NF-κB in both KO-sensitive and -resistant than ART and RIF (Figure 9F2).

Thirdly, PXR in livers from KO-sensitive or -resistant under CMC-Na treatment declined by contrast with WT-sensitive or -resistant, respectively (Figure 9G1). ART decreased it in KO-sensitive but remarkably enhanced it in KO-resistant. ART-CHR also led to a higher hepatic and enteric PXR in KO-sensitive or-resistant. In Figure 9H1, KO-resistant showed an activated hepatic CAR than WT-resistant no matter whether ART administration or not. NF-κB in livers from sensitive parasites infected mice was repressed by knocking out of Mdr1a (Figure 9I1). ART gave a minor augmented NF-κB level in KO-sensitive when compared with WT-sensitive; it was also enhanced by ART-CHR in contrast with WT- and KO-sensitive or WT- and KO-resistant. As for intestinal PXR mRNA, it was activated after knocking out Mdr1a gene (Figure 9G2) in both sensitive and resistant groups. Similar results were observed under dealing with ART alone or combination. In Figure 9H2, enteral CAR mRNA got higher in KO-sensitive mice than WT-sensitive mice before and after ART administration. However, ART-CHR combination decreased it in both KO-sensitive and -resistant mice. Deficiency of Mdr1a significantly up-regulated intestinal NF-κB mRNA pre- and post-administration of ART-CHR (Figure 9I2).

## DISCUSSION

To date, there have been of considerable interests among basic experimental and clinical researchers attempting to elucidate the resistance mechanisms (18-23). Our results also experimentally demonstrated the potential of ART resistance due to a slightly downed antimalarial activity against its resistant *P. berghei*. Delayed or slow growth cycle in RBC for resistant parasites was discovered that might be helpful to explain how they always could escape from the chemotherapy, now that ART is most efficient to kill those developed in trophozoit and schizont stages.

Secondly, this study proved that ART resistance is likely interrelated to some crucial members, such as P-gp, Bcrp, and MRPs. It was commonly thought that currently used ART derivatives as non-P-gp substrates are not transported by P-gp (24). Theoretically, P-gp-mediated MDR thus should not be the critical mechanism of ART itself resistance, yet ART as P-gp inducer has been reported to slightly induce P-gp-mediated (N)-methyl-quinidine (NMQ) transport in HEK293 (25). ART increased net secretion of digoxin, P-gp probe substrate, in human intestinal epithelial cells (T84) (26). Currently, we found that ART up-regulated P-gp/Mdr1 mRNA in small intestines after being infected with sensitive and resistant parasites, which might result in an enhanced transmembrane efflux of P-gp substrate partners in ACTs and leads to MDR. Concurrently, CHR has the potential to re-sensitize the parasites towards ACTs by blocking P-gp/Mdr1 mRNA expression levels. It is worthy of note that CHR itself as well as VER killed resistant parasites notwithstanding less so in the parental sensitive groups and NOV treatment. The underlying mechanism how it works is not clear now.

Another intriguing thing is that although ABCG2 mRNA were elevated as well as Mdr1 mRNA by ART and dropped by CHR or ART-CHR, its protein levels were oppositely regulated, which was in accordance with the results reported in our previous studies and the literature (16, 25). ABCG2 functions as the transporter of porphyrins such as heme and defends cells and/or tissues from protoporphyrin accumulation in response to hypoxia. For this reason, ABCG2 expression is always up-regulated in tissues exposed to low-oxygen environments. Parasite invasion always leads to focal ischemia and hypoxia by rupturing the RBC of host. ABCG2 thus might play an indefinite but crucial role in ART resistance mechanism that requires to be further studied.

Additionally, MRP1/ABCC1, widely expressed in various tissues including liver, intestine, macrophages, and RBC (27), seems to play a critical modulation role in cellular oxidatitive stress and redox homeostasis. Despite distant evolutionary distance between MRP1 and P-gp, gene transfer tests hinted that they might confer resistance to a similar, even not identical, range of drugs. Our data demonstrated that ART dropped enteric MRP1 mRNA expressions in healthy or sensitive mice but instead facilitated it when mice were infected with resistant parasites. CHR significantly suppressed MRP1 mRNA expression in healthy, sensitive, and resistant parasites infected mice intestines and ART-CHR combination could effectively restrain ART-induced MRP1 mRNA upregulation. MRP1, therefore, potentially functions as another critical member of ABC transporter for CHR to reverse ART resistance. Furthermore, this current study suggested that MRP2, MRP4, or MRP5 might also to some extent be involved in the reversing mechanism for CHR. Regretfully, how CHR modulates these subfamily transporters especially the interactions between them has not been understood well yet.

PXR and CAR, so called “low affinity, high capacity” receptors, belongs to orphan nuclear receptors abundantly expressed in the liver, intestine, and kidneys. Both of them modulate some important ABC transporters and process highly overlapped target genes whereas they share much less similarities in their activation mechanisms. PXR activation is exclusively dependent on ligand-binding, while CAR could be activated by either direct ligand binding or ligand independent (indirect) pathways. We discovered that ART did not change transcriptional levels of PXR or CAR using LS174T cell lines, whereas AE slightly descended PXR versus NC, which is not in accordance with the literature (5). CHR and ART-CHR activated *in vitro* transcriptional expression of PXR/CAR. But *in vivo* hepatic and enteric PXR/CAR and NF-κB p52 mRNA in Mdr1a WT- or KO-sensitive and -resistant indicated complex. Invasion of parasites in Mdr1a+/+ mice led to opposite expressions of PXR/CAR and NF-κB not only between livers and small intestines but also between sensitive and resistant *P. berghei*. Likewise, ART-CHR also exhibited reversed modulation on PXR/CAR mRNA in livers and small intestines; however, it caused consistent down-regulation on hepatic and enteric NF-κB positively correlated with P-gp expression (28). It hints that NF-κB signaling pathway might play a crucial role for CHR in depressing Mdr1 expression.

PXR/CAR and NF-κB p52 in Mdr1a-/- mice were also detected in this paper. We observed that infection of resistant parasites were inclined to increase them versus sensitives except declined enteric CAR and hepatic NF-κB by ART-CHR. When compared with WT and KO mice, hepatic PXR/CAR and NF-κB were reversely regulated relative to those in small intestines with the exception of enhanced hepatic and enteric PXR by ART-CHR. Intriguingly, ART-CHR boosted hepatic and intestinal NF-κB in KO-sensitive or KO-resistant mice. This repeatedly evidenced that NF-κB should function as a crucial nuclear transcriptional factor for CHR.

Besides, ART-CHR accreted hepatic PXR and intestinal CAR mRNA but depressed hepatic CAR and intestinal PXR in WT-resistant mice versus WT-sensitive mice. In KO-sensitive and -resistant mice, however, both PXR and CAR mRNA levels were boosted by ART-CHR. When compared with WT-resistant and KO-resistant, PXR in both livers and small intestines was increased whereas intestinal CAR mRNA declined. This fickle function on PXR/CAR by CHR requires to be further clarified.

In conclusion, we found that CHR might be a promising candidate inhibitor of ART resistance. PXR/CAR and NF-κB signaling pathway potentially involved in the mechanism. This paper thus provided a good policy to solve the bottleneck problem in ACTs clinical therapy.

## METHODS

### Chemicals and reagents

ART (colourless needle crystal) was purchased from Chongqing Huali Konggu Co., Ltd (CAS: 6398-64-9, purity ≥99.0%, Chongqing, China). CHR (yellow crystal with purity ≥98.0%) was purified in our lab from the acetone layer of ART industrial waste materials by using multiple column chromatography methods as described previously (14). The acetone layer of ART industrial wastes were kindly provided by Chongqing Huali Konggu Co., Ltd in 2010 and the voucher specimen (20100102) has been currently deposited with Medical College, Yangzhou University, for further references. Verapamil hydrochloride tablets (VER), well-known as a specific P-gp inhibitor (29-31), were purchased from Guangdong Huanan Pharmaceutical Group Co., Ltd (No.140101, Guangdong, China). Novobiocin (NOV), as a specific Bcrp inhibitor (32), was purchased from Hefei Bomei Biotechnology Co., Ltd (CAS: 1476-53-5, purity ≥90%, Hefei, China). Artesunate (AE, purity>98%), one of ART analogues, was purchased from Sigma (Sigma, USA). Rifampicin (RIF) and 5-fluorouracil (5-FU), being as PXR/CAR receptor agonist, were obtained from Beijing Solarbio Science and Technology Co. Ltd.

Sodium carboxymethyl cellulose (CMC-Na) was purchased from Tianjin Guangfu Fine Chemical Research Institute (Tianjin, China). Sodium citrate was purchased from Tianjin Damao Chemical Research Factory (Tianjin, China). Absolute ethyl alcohol, isopropanol, and trichlormethane were purchased from Sinopharm Chemical Reagent Co. Ltd. All the reagents were of analytical grade.

### Recovery and passage of ART-sensitive or -resistant *P. berghei* K173

ART-sensitive *P. berghei* K173 was from National Engineering and Technology Research Center for the Development of New Chinese Compound Medicines (Beijing, China). Resistant parasites were obtained by passaging the sensitives under incremental ART dosages until 42 generations and kindly provided by Prof. Dai from Chengdu University of Traditional Chinese Medicine (Sichuan, China). The resistance index was 14.15 (33). Both sensitive and resistant parasites were preserved in mice blood at -80°C before use.

Cryopreserved *P. berghei* were completely thawed within 1 min under the temperature of 38ºC and continuous shaking. Then the blood was diluted with sodium citrate-normal saline to promise each 0.2 mL contained approximately 10^6^ parasite infected erythrocytes. Inbred male ICR mice weighing an average of 20 g were respectively infected by intraperitoneal injection (i.p.) with 0.2 mL of parent sensitive or resistant *P. berghei* infected red blood cells (RBC) under a strict sterile manipulation. Blood smear was freshly prepared by Giemsa stain method and then monitored in the oil immersion lens of microscope (40×) to calculate parasitaemia% according to the formula below. Parasitaemia% was traced twice per day aiming to indicate the infection difference between sensitive and resistant parasites within an experimental time period. The mean parasitaemia% was calculated as follows:

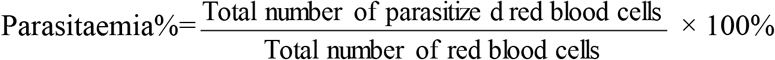

Donor *P. berghei* infected mice with parasitaemia% over 20% were sacrificed and the blood samples were collected from the orbit for later experiments afterwards.

### *In vivo* anti-malarial activity of drugs against ART-sensitive and -resistant *P. berghei*

Healthy male ICR mice, weighing 18-22 g, were purchased from Animal Center of Ningxia Medical University (Ningxia, China). Male inbred Abc1a deficiency (knock-out, KO) mice were purchased from Shanghai Model Organisms (China) with FVB background. All animals within weight range of 18-22 g were housed in polycarbonate cages and acclimated in an environmentally controlled room (23 ± 2ºC, with adequate ventilation and a 12 h light/dark cycle) at least 7 days prior to use. They were maintained on standard pellet diet and water *ad libitum* before and during the experiments. The protocol was submitted to and approved by the University Ethics Committee (20170120). All procedures involving animals were in accordance with the Regulations of the Experimental Animal Administration, State Committee of Science and Technology.

The animals were randomly assigned to three groups (*n*=60 for each group) including uninfected (healthy) and ART-sensitive or -resistant *P. berghei* infected groups. On day 0 (before starting oral administration of drugs), mice were intraperitoneally inoculated with 0.2 mL of infected blood containing about 10^6^ parasitized RBC, which is expected to produce steadily rising consistent infection of the required intensity (when parasitaemia% reached at least 5%) in mice. Each group was thereafter split into five groups (*n*=12 for each group) including negative control (0.5% sodium carboxymethylcellulose, CMC-Na), NOV or VER alone (positive control, 100 mg/kg), ART alone (40 mg/kg), ART-CHR combination (1:2, 40:80 mg/kg), and CHR alone (80 mg/kg). The drugs were administrated intragastrically once per day for consecutive seven days, respectively. The body weights and survivals of mice were recorded once each day. At day 7, one hour after drug administration, the mice were sacrificed by cervical vertebra dislocation under anesthesia. Ileum and jejunum were harvested and cleaned using normal saline at least three times and stored at -80ºC before use.

The mean inhibition% was calculated as follows:

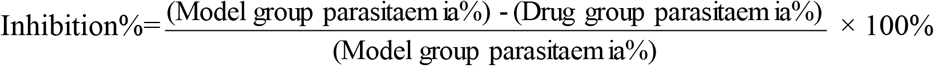

### ABC transporter mRNA expressions detected by real-time quantitative polymerase chain reaction (RT-qPCR)

Total RNA was extracted from the mouse small intestine using E.Z.N.A.™ Total RNA Kit (OMEGA bio-tek, Norcross, GA, USA) in accordance with the manufacturer’s instructions. Its concentration was measured with a microplate spectrophotometer (Bio-RAD, USA) at a wavelength of 260 nm. The quality was assessed by agarose gel electrophoresis method. Qualified total RNA (3 μg) was thereafter reversely transcribed into first strand complementary DNA (cDNA) using Thermo Scientific RevertAid Fisrt Strand cDNA Synthesis Kit (Thermo Scientific). Each cDNA sample (1 μL) was amplified with 10 μL of Thermo Scientific Maxima SYBR Green qPCR Master Mix (2×), ROX Solution (Thermo Scientific), and 1 μM of each primer. Amplification was performed in a RT-qPCR IQ5 System (Applied Biosystems, Foster City USA) with the following parameters: denaturation at 94ºC for 30 s followed by 45 cycles of denaturation at 94ºC for 30 s, annealing at 60ºC for 30 s, and extension at 72ºC for 10 s. End extension time was 2 min.

The sequences of the oligonucleotide primers for genes of ABC transporters used in this study were elaborately displayed in Table 2. The relative mRNA expression levels of these ABC transporters in each sample (normalized to that of GAPDH) were determined using 2^-ΔΔCt^ method. All RT-qPCR experiments were repeated three times.

**TABLE 2.**
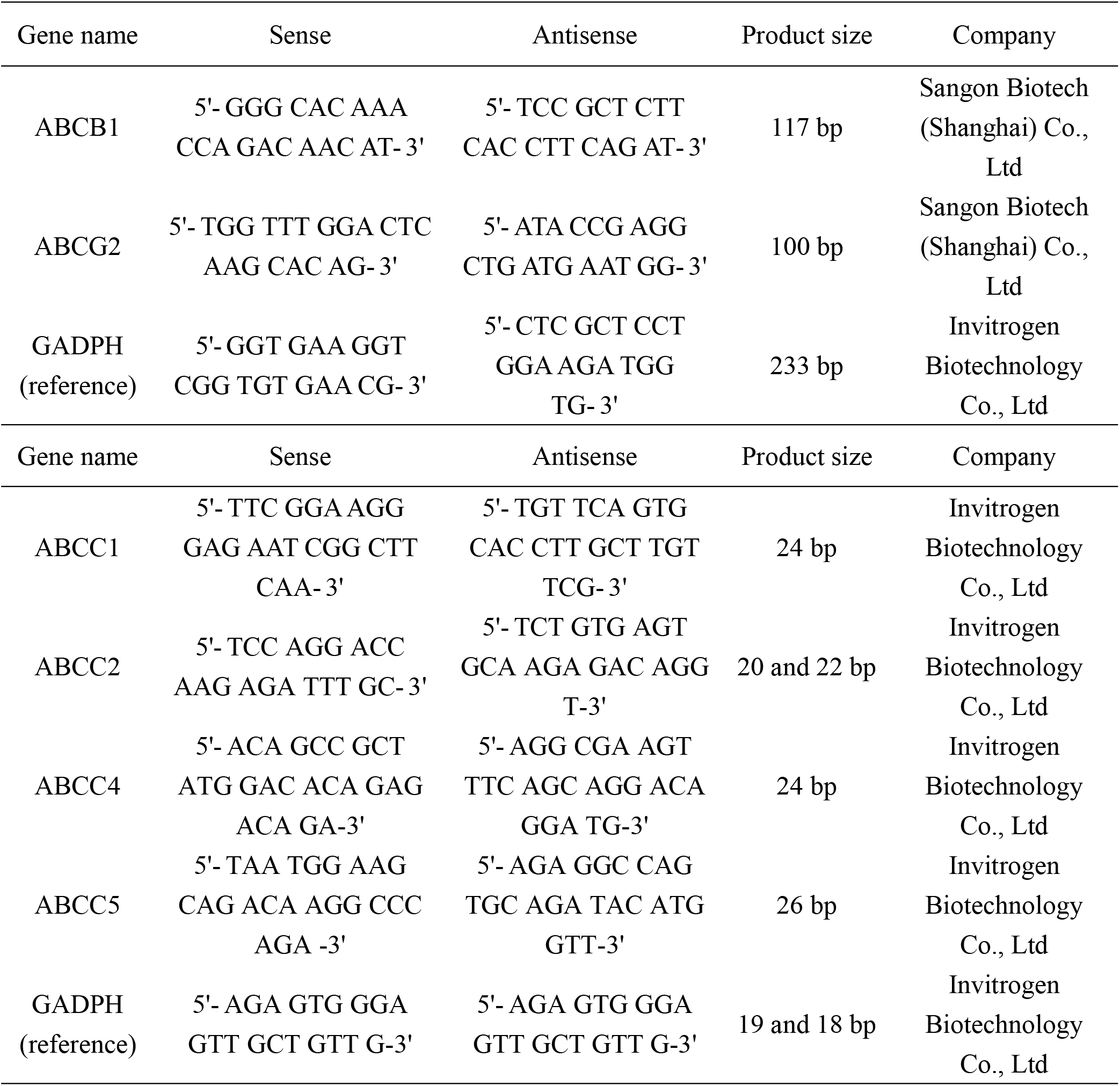
The sequences of the oligonucleotide primers for genes of ABC transporters.

### P-gp/Bcrp protein levels in small intestines detected by western-blot analysis

The total membrane proteins were harvested by KenGEN Whole Cell Lysis Assay Kit (KenGEN BioTECH, Nanjing, China), and the protein concentrations were assayed using KeyGEN BCA Protein Quantitation Assay kit (Nanjing KeyGEN Biotech, China). An equal quantity of protein (80 μg) from total protein was resolved using 7.5% SDS-PAGE gel and subsequently transferred onto nitrocellulose membranes (Bio-Trace). After blocking the membrane with 5% non-fat milk in Tris-buffered saline (Biotopped) at room temperature for 1 h, the membrane was incubated at 4ºC for 12 h with rabbit monoclonal primary antibodies against Mdr1a (1:1250; ab170904) and Anti-Sodium Potassium ATPase antibody-Plasma Membrane Loading Control (1:20000; ab76020) or rabbit polyclonal antibody against ABCG2 (1:100; sc-25822, Santa Cruz) and mouse monoclonal antibody against *β*-actin (1:150; sc-47778, Santa Cruz). Antibodies were purchased from Abcam (Cambridge, United Kingdom). The membranes were incubated with horseradish Peroxidase-conjugated AffiniPure Goat Anti-Rabbit IgG and Goat Anti-Mouse IgG (ZSGB-BIO, Beijing, China) for 2 h and signals were observed using Super Signal West Pico (Thermo Scientific). Western blotting band intensity was quantified by densitometry analysis using ImageJ version 2x (NIH Image software, Bethesda, MA, USA).

### Transcription level of PXR and CAR using a dual-luciferase reporter assay in the human intestinal carcino cell lines LS174T

The human colon adenocarcinoma cell lines LS174T (derived from Caucasian colon adenocarcinoma) maintained in RPMI 1640 medium (Gibco) supplemented with 10% FBS (BI) were purchased from Tongpai (Shanghai) Biotechnology Co., Ltd and cultured at 37ºC under a humidified atmosphere of 5% CO_2_. CCK8 cell proliferation-cytotoxicity detection kit was used to determine IC_50_ values of different drugs including 5-FU, RIF, ART, CHR, and CHR-ART combination (2:1). Dual-luciferase reporter assay was utilized to detect the transcription levels of hPXR and hCAR receptors, using RIF and 5-FU as positive controls. 0.1% DMSO was used as a negative control. The reporter gene plasmids encoding MDR1, human PXR and CAR were constructed according to the literature (5) and made some necessary modification. The reporter gene plasmid for MDR1 was constructed (Fig. 8A) by cloning and ligating the corresponding *XhoI* fragment of the MDR1 promoter (−1830 to +281, -7975 to -7013), into the *XhoI* site of PGMLV-rev vector (Jiman Biotechnology Co., Ltd, Shanghai, China). The MDR1 downstream promoter region encompassing the sequence between bases -1830 and +281 was amplified by PCR out of human genomic DNA using the primers as shown in Table 3, which introduce *MIuI* and *Bsal* sites, respectively. The PCR fragment digested with *MIuI* and *Bsal* was ligated into *MIuI*/*Bsal* digested PGMLV-rev vector, creating p-1830, and sequenced. The MDR1 promoter fragment (−7975 to -7013) containing the cluster of nuclear receptor response elements was amplified by PCR with the primers (Table 3) which introduce *Bsal* and *SalI* sites, respectively. To construct a human PXR or CAR expressed plasmid, the open reading frame of human PXR/CAR was amplified by PCR using the primers (Table 3) which introduced *XhoI* and *BamhI* sites, and sequenced.

**TABLE 3.**
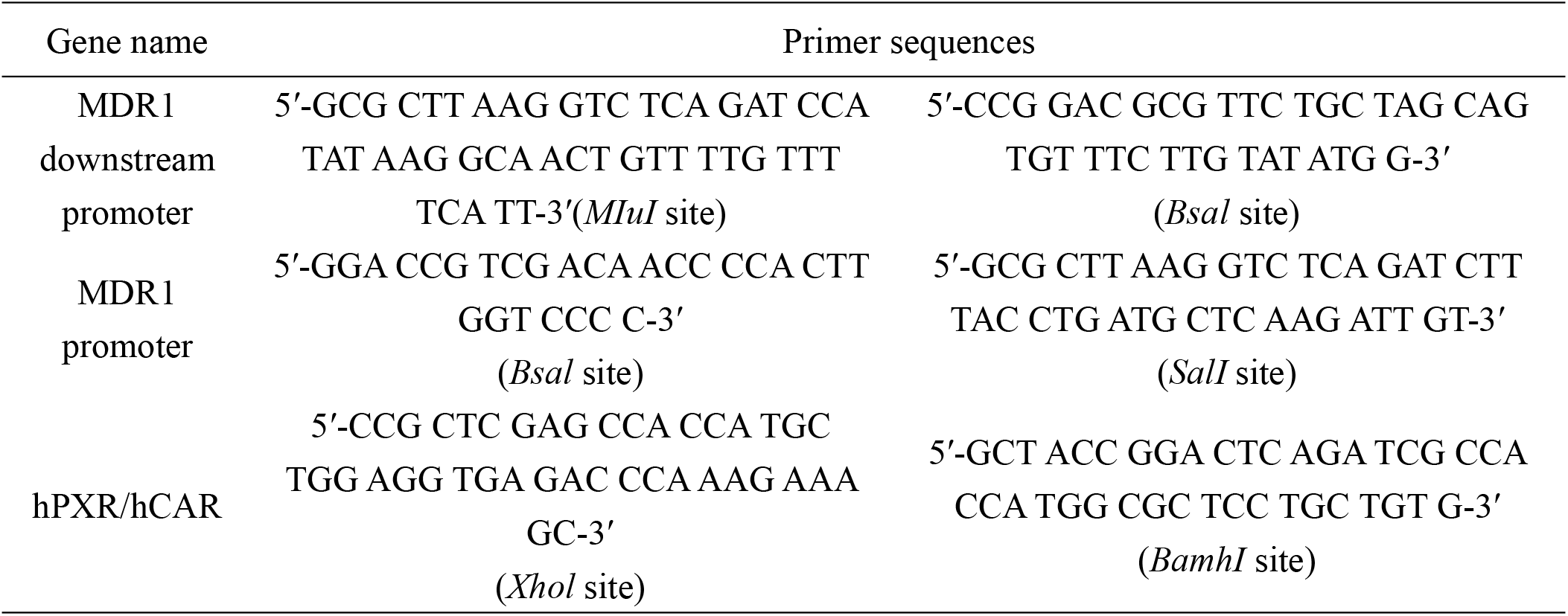
The primers of reporter gene plasmid for MDR1 as well as hPXR/CAR.

LS174T cells were cultivated in 24-well plates at a density of 1.5 × 10^5^ cells/well per day before transfection. The plasmid DNA was transfected by using Lipofectamine™ 2000 transfection reagent (ThermoFisher Scientific Inc.). In brief, 238 ng of gene over-expressed plasmid (OE/NC) and 238 ng of promotor reporter gene plasmid, 24 ng of pRL-TK, 50 μL of serum-free DMEM, and 1.5 μL of Lipofectamine™ 2000 transfection reagent was added into the 96-wells. After 24-hour incubation, the cells were washed with phosphate-buffered saline and lysed with 100 μL of 5x passive Lysis Buffer. After centrifugation, cleared lysates were respectively stored at -80ºC in the eppendorf (EP) tubes before use for reporter gene assays. Transfections were done in duplicate, and experiments were repeated three times. Drugs including ART, ARE, CHR, RIF, and 5-FU at optimized dosage were used to treat the lysates separately. Luciferase measurements were done according to the manufacturer’s recommendations. To identify statistically significant differences, statistical tests were done as indicated with the mean values of at least three independent experiments done in triplicates using Graphpad Prism 7.0 (USA).

### *In vivo* PXR/CAR and NF-κB p52 mRNA expression in livers and small intestines from Abcb1 WT- or KO-sensitive or -resistant mice

Gene primers of PXR/CAR and NF-κB were provided by Beijing Robby Biological Technology Co. Ltd. HiScript III RT SuperMix for qPCR (+gDNA wiper) and ChamQ Universal SYBR qPCR Master were purchased from Nanjing Novizan Biotechnology Co. Ltd. Trizol, agarose, NA-red, DNA ladder, DEPC treated water, TAE electrophoresis buffer, and DNA loading buffer were purchased from Beyotime Biotechnology. The sequences of the oligonucleotide primers for genes of PXR/CAR and NF-κB p52 used in this study were listed in Table 4. Detailed RT-qPCR semiqunantitative method referred to that as mentioned in mRNA expression of ABC transporters. In brief, amplification was performed in a RT-qPCR IQ5 System (Applied Biosystems, Foster City USA) with the following parameters: denaturation at 95ºC for 10 m followed by 40 cycles of denaturation at 95ºC for 15 s, annealing and extension at 60ºC for 60 s.The relative mRNA expression levels of PXR/CAR and NF-κB p52 in each sample (normalized to that of β-actin) were determined using 2^-ΔΔCt^ method. All RT-qPCR experiments were repeated three times.

Differed PXR/CAR as well as NF-κB p52 mRNA levels, between WT and Mdr1a deficiency mice after being infected with sensitive or resistant malarial parasites under different drug treatments, were therefore compared, respectively.

**TABLE 4.**
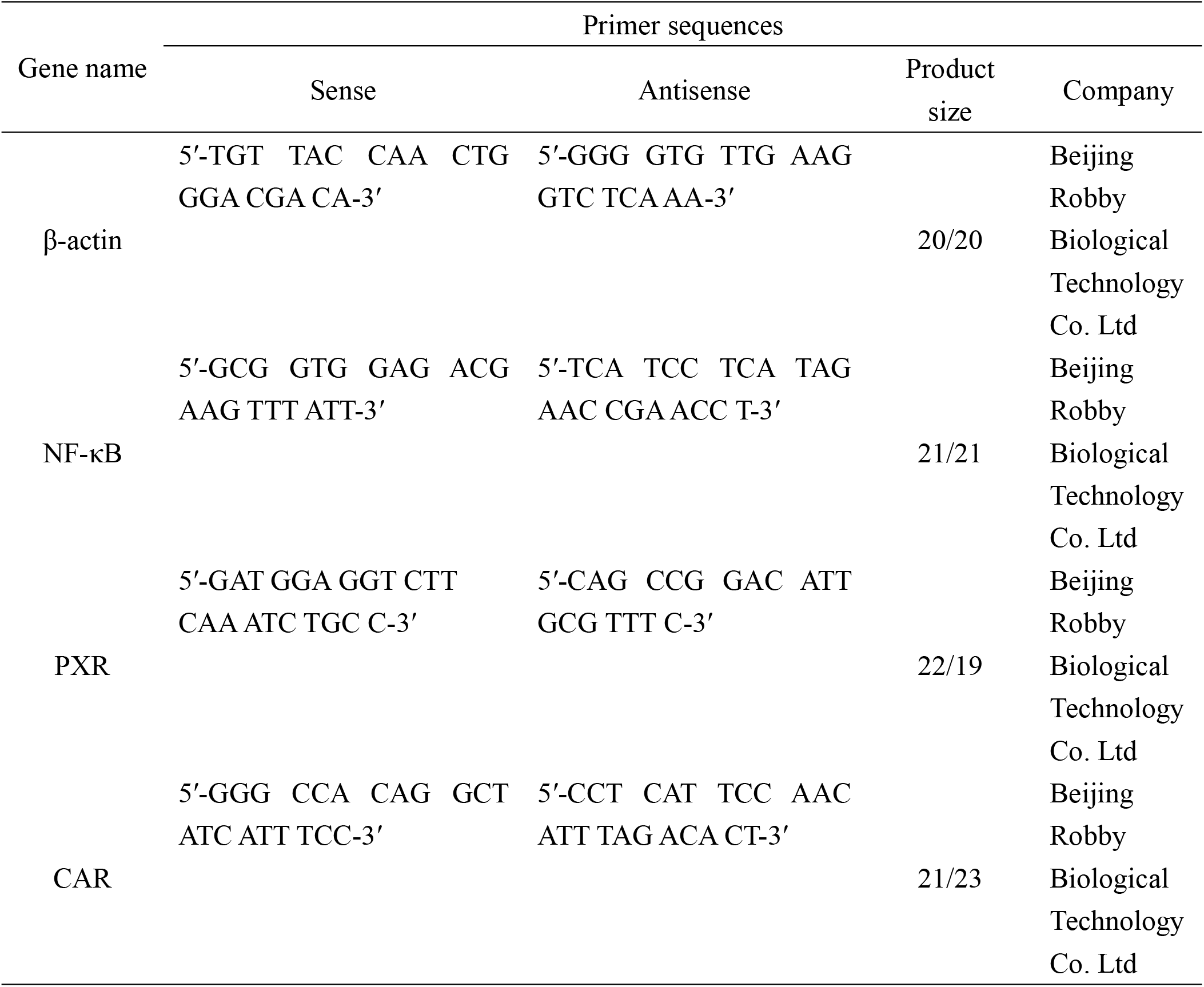
The primers of PXR/CAR and NF-κB p52.

### Statistical Analysis

Survival curves are estimated using the Kaplan-Meier method and compared statistically using Log-rank (Mantel-Cox) test (Graphpad Prism 7.0, USA).

Other data was analyzed using the SPSS 18.0 software (IBM, USA) or Graphpad Prism 7.0 and submitted to a one-way analysis of variance (ANOVA) to detect significant differences between study groups. Turkey’s test was applied to identify any difference between means at a significance level of *P*<0.05 or *P*<0.01.

## Abbreviations

MDR: multi-drug resistance
ABC transporters: ATP binding cassette transporters
P-gp: P-glycoprotein
Bcrp: breast cancer resistance protein
MRPs: multi-drug resistance proteins
ACTs: artemisinin based combination treatments
ART: artemisinin
CHR: chrysosplenetin
5-FU: 5-fluorouracil
RFP or RIF: rifampin
NOV: novobiocin
VER: verapamil hydrochloride
CMC-Na: sodium carboxymethyl cellulose
CCK8: cell counting kit
NC: normal control
DMSO: dimethyl sulfoxide
RBC: red blood cell
PXR: pregnane X receptor
CAR: constitutive androstane receptor
NF-κB: nuclear factor kappa-B

## ACKNOWLEGEMENTS

This work was financially supported by the National Natural Science Foundation of China (No. 81760377), Science and Technology Funds of Nanjing Medical University (NMUB2019275), and Science and Technology Funds of Kangda College, Nanjing Medical University (KD2019KYJJZD003). The funding sources did not play any role in the design, conduct or interpretation of our outcomes.

